# Simultaneous analysis of antigen-specific B and T cells after SARS-CoV-2 infection and vaccination

**DOI:** 10.1101/2021.12.08.471684

**Authors:** Krista L. Newell, Mitchell J. Waldran, Stephen J. Thomas, Timothy P. Endy, Adam T. Waickman

## Abstract

Conventional methods for quantifying and phenotyping antigen-specific lymphocytes can rapidly deplete irreplaceable specimens. This is due to the fact that antigen-specific T and B cells have historically been analyzed in independent assays each requiring millions of cells. A technique that facilitates the simultaneous detection of antigen-specific T and B cells would allow for more thorough immune profiling with significantly reduced sample requirements. To this end, we developed the B And T cell Tandem Lymphocyte Evaluation (BATTLE) assay, which allows for the simultaneous identification of SARS-CoV-2 Spike reactive T and B cells using an optimized Activation Induced Marker (AIM) T cell assay and dual-color B cell antigen probes. Using this assay, we demonstrate that antigen-specific B and T cell subsets can be identified simultaneously using conventional flow cytometry platforms and provide insight into the differential effects of mRNA vaccination on B and T cell populations following natural SARS-CoV-2 infection.

## Introduction

The identification, quantification, and characterization of antigen-specific lymphocytes is a critical prerequisite for understanding the breadth, magnitude, and functional potential of vaccine and/or natural pathogen-elicited immunity. Multiple distinct methodologic approaches have been developed that allow for the identification of antigen-specific T and B cells. For the identification of antigen-specific T cells, these include quantification of functional cytokine-producing T cells by ELISPOT, flow cytometric detection of intracellular cytokine production following peptide stimulation, staining with peptide/HLA probes, and upregulation of activation-induced markers following antigen engagement (Altman et al., 1996; Kern et al., 2000; Maecker et al., 2001; Frentsch et al., 2005; Dan et al., 2016, 2021). Analogous approaches have also been developed for the identification of antigen specific B cells including, but not limited to, B cell ELISPOT assays and the use of fluorescently conjugated protein probes to stain antigen-specific B cells (Julius et al., 1972; Czerkinsky et al., 1983; Moody and Haynes, 2008; Boonyaratanakornkit and Taylor, 2019). However, a major limitation of the approaches developed thus far is that they interrogate rare B and T cell subsets in isolation. An approach capable of identifying and phenotyping antigen-specific B cells, CD4^+^ T cells, and CD8^+^ T cells simultaneously would not only preserve valuable samples by performing multiple assays in parallel, but would also permit the efficient longitudinal assessment of B and T cells that recognize the same antigen over time as they expand and contract following challenge.

In this study, we endeavored to simultaneously identify and characterize SARS-CoV-2 spike protein-specific B cells, CD4^+^ T cells, and CD8^+^ T cells in the same PBMC sample. We used the combination of an optimized Activation-Induced Marker (AIM) assay to identify spike reactive T cells following stimulation with overlapping spike peptide pools, and spike reactive B cells by baiting with dual spike trimer fluorescent tetramers. This approach was then used to quantify and longitudinally track the T and B cell populations induced by natural SARS-CoV-2 infection and subsequent SARS-CoV-2 mRNA vaccination in the same individuals. Flow cytometric analysis of these antigen-specific populations revealed that while the spike-specific B cell, and to lesser degree CD8^+^ T cell responses, were consistently enhanced by mRNA vaccination, a clear boosting effect on spike-specific CD4^+^ T cells was only detectable in individuals with low or undetectable T cell responses following natural infection. Furthermore, we validated our observations of SARS-CoV-2-specific B cell functional enhancement upon vaccination with paired serological analysis. Our findings suggest divergent patterns of B and T cell subset mobilization with repeated antigen encounter and highlight the utility of examining these populations in concert.

## Materials and Methods

### Study design

PBMC and sera for this study were obtained from the previously described SUNY Upstate Medical University Convalescent COVID-19 Plasma Donor protocol (Fang et al., 2021; Newell et al., 2021). A subset of 10 donors was selected for focused analysis in the study using the following criteria: 1) blood collection at least 30 days after but not longer than 75 days after their last COVID-19 symptom, 2) immunization with two doses of a SARS-CoV-2 mRNA vaccine at least 9 months after a negative PCR test (8/10 Pfizer-BioNTech, 2/10 Moderna mRNA-1273), and 3) blood collection at least 30 days, but not longer than 22 weeks following vaccination. One subject was excluded from the final analysis due to immunosuppressive treatment for autoimmune disease at the time of sample collection. Archived sera and PBMC from healthy individuals collected prior to December 2019 were used as negative controls. Sample size was determined based on subject availability and conformity of sample collection to the described timeline.

### Spike tetramer generation

Full-length SARS-CoV-2 Spike (2P-stabilized, C-terminal Histidine/Avi-tagged) was obtained from BEI resources (Manassas, VA, USA, Cat. NR53524) and biotinylated using a BirA ubiquitin ligase (Avidity, Aurora, CO, USA, Cat. Bir500A) following the manufacturer’s recommended protocol. Biotinylated spike protein was purified using a 40K MWCO 2 mL Pierce Zeba™ desalting column (Thermo Fisher Scientific, Waltham, MA, USA, Cat. 87768), and mixed with streptavidin BV421 (BD, Cat. 563259) or streptavidin allophycocyanin (APC) (BD Cat. 554067) separately at a 20:1 ratio (~6:1 molar ratio) as previously described (Dan et al., 2021). Tetramerized Spike probes were stored at 4°C until use. Just prior to probe staining, spike protein probes were added one-by-one to FACS wash buffer (1x PBS, 2% fetal bovine serum) containing 5μM free d-biotin (Avidity, Cat. Bir500A). Both streptavidin-fluor conjugates were used to stain DMSO control samples to further verify the absence of significant frequencies of non-specific streptavidin-binding B cells.

### Activation Induced Marker (AIM) assay

Spike-specific T cells were quantitated as a frequency of AIM^+^ (OX40^+^CD69^+^) CD4^+^ T or (CD25^+^CD69^+^) CD8^+^ T cells among PBMC following 24 hours of stimulation with overlapping peptide pools covering the length of SARS-CoV-2 spike peptide (BEI, Cat. NR52402), as previously described (Fang et al., 2021). Cryopreserved PBMC were thawed, washed twice, and cultured in complete cell culture media: RPMI 1640 medium (Corning, Tewksbury, MA, USA, Cat. 10-040-CV) supplemented with 10% heat-inactivated fetal bovine serum (Corning, Cat. 35-010-CV), 1% L-glutamine (Lonza, Basel, Switzerland, Cat. BW17-605E), and 1% Penicillin/Streptomycin (Gibco, Waltham, MA, USA, Cat. 15140122). Cellular viability was assessed by trypan blue exclusion and cells were resuspended at a concentration of 5 × 10^6^ cells/mL. Next, spike peptide pool cocktail was added to each well, diluted to a final concentration of 1 μg/mL/peptide (DMSO concentration 0.5%). Incubation with an equimolar amount of DMSO (0.5%) was used as a negative control, or monoclonal anti-human CD3 (Mabtech, Sweden, Cat. 3420-2AST-10) at a 1:1000 dilution for positive control wells. Cells were cultured for 24 hours at 37°C, 5% CO_2_ in polystyrene plates.

### Staining and Flow Cytometry

Stimulated PBMC were washed twice in warm complete culture media and transferred to polypropylene 96-well plates for staining in the dark with 50μL of spike tetramer probe cocktail containing 100ng Spike per probe (total 200ng) at 4°C for 1 hour prior to surface staining in FACS wash buffer with the antibodies listed in table S1 at 4°C for 30min. Cells were washed with 1x PBS to avoid binding of antibodies to viability dye and stained with the LIVE/DEAD Fixable Aqua Stain Kit (Thermo, Cat. L34957) in 1x PBS at 4°C for 30min to exclude dead cells. The cells were washed again with FACS buffer to quench remaining dye. Single-color compensation controls were prepared using Onecomp eBeads™ compensation beads (Thermo, Cat. 01111142) and Arc™ Amine Reactive Compensation Bead Kit (Thermo, Cat. A10628) according to manufacturer’s instructions. For streptavidin-fluorophore compensation, biotin anti-human CD19 (BioLegend, San Diego, CA, USA, Cat. 302203) was used as a primary stain. Stained cells and compensation controls were acquired on a BD Fortessa or Aria II and analyzed using FlowJo (BD), version 10.8.0. The antibody panel utilized in all experiments is shown in table S2. Validation was performed between batches of spike tetramer probes on the same sample to confirm consistency of binding and fluorescence properties.

The frequency of spike-specific B cells was expressed as a percentage of total B cells (CD19^+^, CD3^/^CD14/CD56^/^LIVE/DEAD^-^, lymphocytes), or as number per 10^6^ PBMC. Spike-specific CD4^+^ and CD8^+^ T cell quantities were calculated as background (DMSO) subtracted data.

### ELISA

SARS-CoV-2 Spike/RBD and nucleocapsid antibody titers were quantified using a sandwich ELISA protocol. In brief, 96 well NUNC MaxSorb flat-bottom plates were coated with 3.99 μg/ml recombinant SARS-CoV-2 Wuhan-Hu-1 spike RBD protein (Sino Biological, Beijing, PR China, Cat. 40592-V08B), 1 μg/ml SARS-CoV-2 Wuhan-Hu-1 nucleocapsid protein (Sino, Cat. 40588-V08B), 0.15 μg/mL SARS-CoV-2 alpha variant spike RBD protein (BEI, Cat. NR-54004), or 0.15 μg/mL beta variant spike RBD protein (BEI, Cat. NR-55278) diluted in sterile 1x PBS. Plates were washed and blocked for 30 min. at RT with 0.25% BSA + 1% Normal Goat Serum in 0.1% PBST after overnight incubation 4°C. Serum samples were heat-inactivated for 30 min. at 56°C and serially diluted 4-fold eight times starting at 1:200 prior to incubation for 2 hrs. at RT on the blocked plates. Plates were washed and antigen-specific antibody binding was detected using anti-human IgG HRP (MilliporeSigma, St. Louis, MO, USA, Cat. SAB3701362), or anti-human IgM HRP (SeraCare, Milford, MA, USA, Cat. 5220-0328). Secondary antibody binding was quantified using the Pierce TMB Substrate Kit (Thermo, Cat. 34021) and Synergy HT plate reader (BioTek, Winooski, VT). Antibody binding data were analyzed by nonlinear regression (One site specific binding with Hill slope) to determine EC50 titers, reported as Kd values, in GraphPad Prism 9.1.0 (GraphPad Software, La Jolla, CA). End-point titers of SARS-CoV-2 antigen-reactive serum samples were determined as the reciprocal of the final dilution at which the optical density (OD) was greater than 2× of control SARS-CoV-2 naïve serum.

### Statistical analyses

Statistical analyses were performed using Prism version 9.1.0 (GraphPad), except for correlations. Spearman’s rho rank correlation coefficients were computed in R (version 1.3.1093) using the rcorr function in the Hmisc package (version 4.4-1). Statistical significance was represented as shown: ns: p > 0.05, *: p < 0.05, **: p < 0.01, ***: p < 0.001, and ****: p < 0.0001. Individual analyses performed and sample size (n) are indicated in the figure legend. Statistical tests were chosen based on descriptive statistics for the specified dataset. The column in each column plot indicates the arithmetic mean of the dataset.

#### Study approval

All participants provided written informed consent prior to participation in the study, which was performed according to protocols approved by the Institutional Review Board (IRB) of the SUNY Upstate Medical University under IRB number 1587400 for experimental samples and 1839296-1 for pre-pandemic control samples. All clinical investigation was conducted according to Declaration of Helsinki principles.

#### Graphics

Study design and assay workflow schematics were created with BioRender.com.

## Results

### BATTLE assay

We developed an assay workflow (**Figure 1a**) for the simultaneous detection of SARS-Cov-2 spike-specific T and B cells from cryopreserved PBMC. Cryopreserved PBMC were thawed and stimulated for 24 hours with overlapping peptide pools spanning SARS-CoV-2 spike to stimulate SARS-CoV-2 spike protein-reactive T cells, followed by incubation with fluorescent spike trimer tetramers to bait spike-binding B cells. These steps were followed by staining with fluorescent antibodies and viability dye prior to interrogation by flow cytometry or sorting. The gating strategy for flow cytometry is illustrated in **Supplementary Figure 1**. Using this approach, we were able to identify and quantify SARS-CoV-2 spike-reactive B cells (**Figure 1b**) in tandem with conventional activation-induced marker (AIM) T cell stimulation using as few as 3 x 10^6^ PBMC per sample from SARS-CoV-2 mRNA vaccine recipients. Notably, stimulation with overlapping peptide pools did not interfere with the detection of antigen-specific B cells using fluorescent spike B cell probes (**Figure 1c, d**). Our results demonstrate that B And T cell Tandem Lymphocyte Evaluation (BATTLE) is an efficient assay capable of simultaneously quantifying and isolating antigen-specific B and T cells from a single PBMC sample.

**Figure 1:**
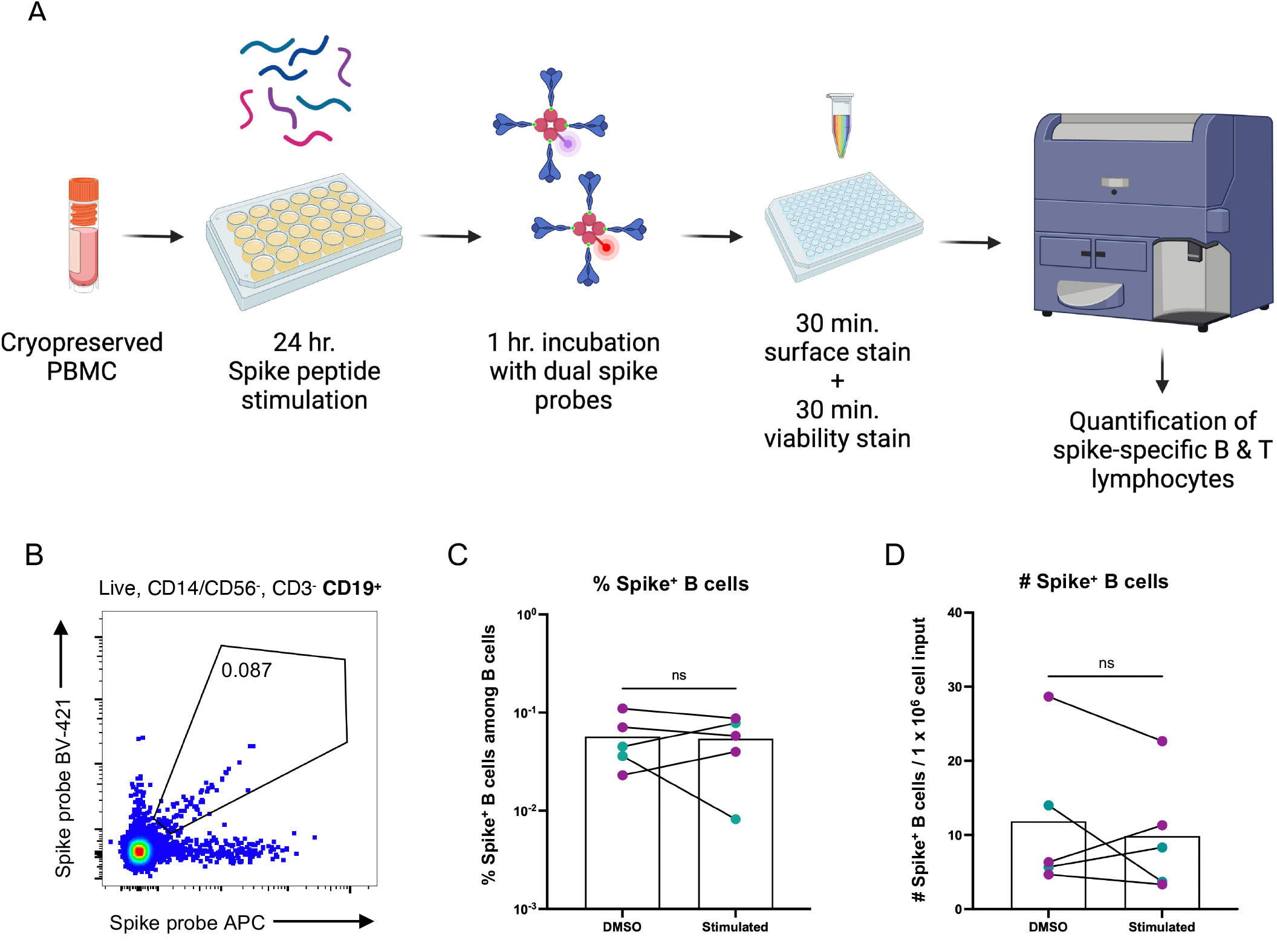
BATTLE: B and T Tandem Lymphocyte Evaluation. A: Schematic of BATTLE assay workflow. B: Representative flow cytometry plot of SARS-CoV-2 spike protein-specific B cells detected from the PBMC of a vaccinated individual using the BATTLE assay. Pre-gating is shown above the plot. The frequency of cells within the double-positive gate among B cells is shown inside the gate. C-D: Total (purple) or monocyte-depleted (teal) PBMC from SARS-CoV-2 mRNA vaccinated individuals were stimulated with overlapping spike peptide pools or DMSO alone and baited with dual spike BCR probes. The frequency among B cells (C) and number per 1 x 10^6^ cells plated (D) of B cells positive for both probes are shown, with each dot representing an individual. Paired DMSO control and peptide pool-stimulated samples from the same individual are connected by a solid line. Paired groups were analyzed by Mann-Whitney U test (n = 5, exact two-tailed p > 0.9999 (C), p = 0.6905 (D)).

### Study design and serology

To demonstrate the utility of our new approach, we used the BATTLE assay to examine the impact of SARS-CoV-2 mRNA vaccination on spike protein-specific adaptive immunity following natural infection. We selected a cohort of subjects from a large study of convalescent plasma donors who experienced symptomatic but mild infection and had a peripheral blood draw between 30 and 75 days of symptom resolution (**Figure 2a**). This window was selected to capture the profile of antigen-specific memory lymphocytes proximally to natural infection, but following resolution of acute inflammatory responses to infection. We included subjects who were vaccinated with two doses of mRNA vaccine at least 9 months following a negative SARS-CoV-2 PCR test and had a subsequent blood draw more than 30 days, but no more than 22 weeks following vaccination. These time constraints were chosen to ensure that immune memory was stable prior to vaccination, and to capture the profile of antigen-specific memory lymphocytes proximally to vaccination, but following resolution of acute vaccine-driven inflammation.

**Figure 2:**
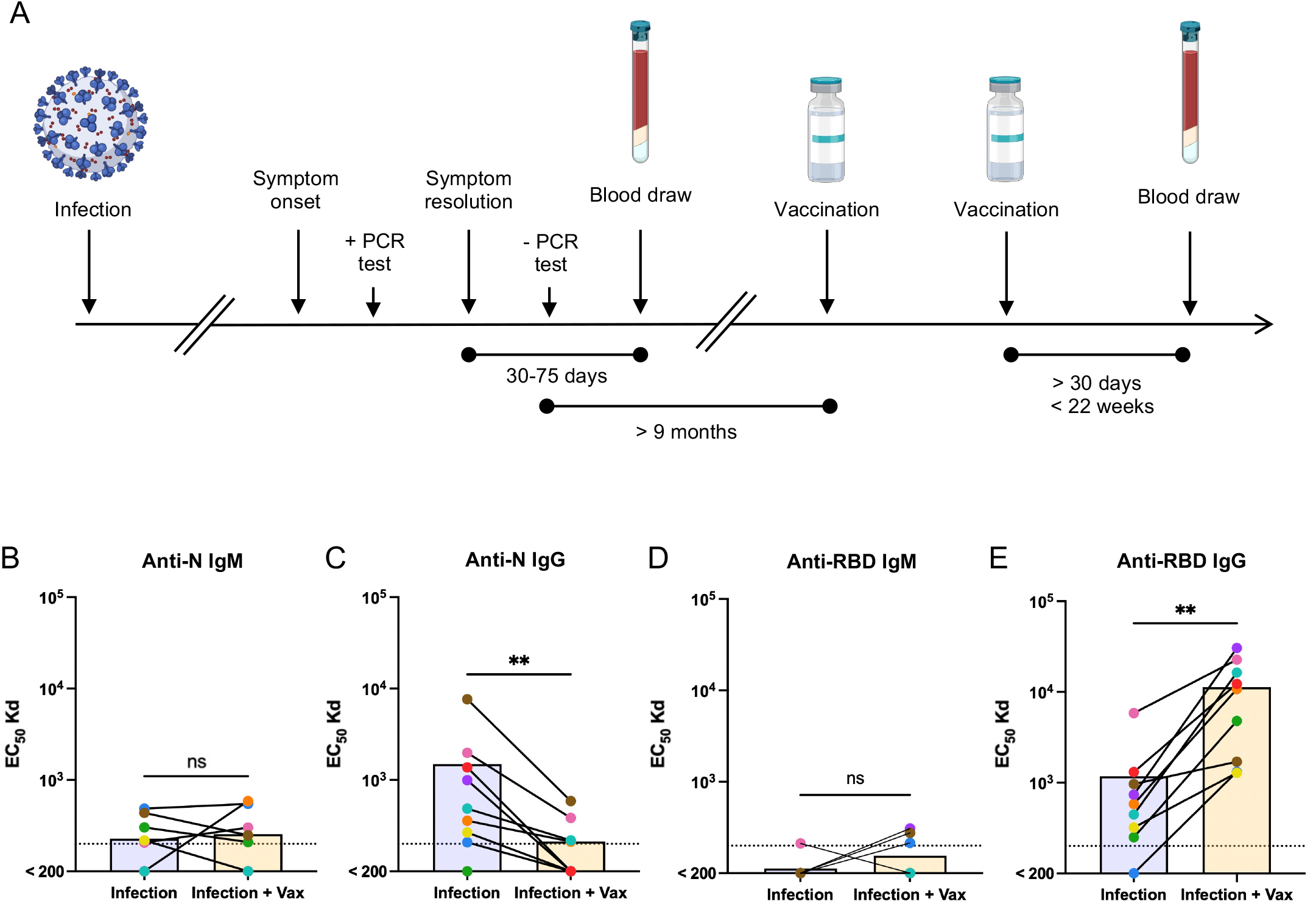
Study design and serology. A: Timeline of SARS-CoV-2 convalescent study and sample collection. B-E: Paired sera following natural infection and after subsequent vaccination from the same individual were assessed for IgM and IgG binding to SARS-CoV-2 nucleocapsid (B, C) or spike RBD (D, E) proteins. Each dot color represents an individual and pairs are connected by a solid line. Differences between infection and infection + vaccination groups were analyzed by Wilcoxon matched-pairs signed rank test (n = 9, exact two-tailed p > 0.9999 (B), p = 0.0078, 0.2500, 0.0039 (C-E)).

To validate and extend the clinical utility of our findings, we evaluated the serological profiles of each donor from the same visits used in the BATTLE assay. As expected, IgG against the SARS-CoV-2 nucleocapsid protein, which is induced by natural infection but not mRNA vaccination, declined between the first and post-vaccination samples, while anti-N IgM was largely below the level of reliable detection (**Figure 2b, c**). Serum anti-spike receptor-binding domain (RBD) IgG, but not IgM, responses also were enhanced by mRNA vaccination (**Figure 2d, e**). Additionally, IgG, but not IgM, raised against the spike receptor binding domain (RBD) from SARS-CoV-2 variant strains alpha and beta was significantly enhanced following mRNA vaccination, although not all subjects had equally improved variant reactivity (**Supplementary Figure 2**).

### SARS-CoV-2 mRNA vaccination significantly increases the number and frequency of spike-reactive B cells among peripheral blood B cells following natural infection

To determine if SARS-CoV-2 mRNA vaccination had an impact on the B cell response targeting spike protein, we examined spike-reactive B cells within post-infection and post-infection plus vaccination PBMC samples. We observed a significant increase in both the frequency and number of spike tetramer double-positive B cells in post-vaccination samples, compared to natural infection alone (**Figure 3a, d, Supplementary Figure 3a**). Importantly, spike-specific B cells were virtually absent in PBMC samples collected prior to the onset of the SARS-CoV-2 pandemic, demonstrating the specificity of the assay.

**Figure 3:**
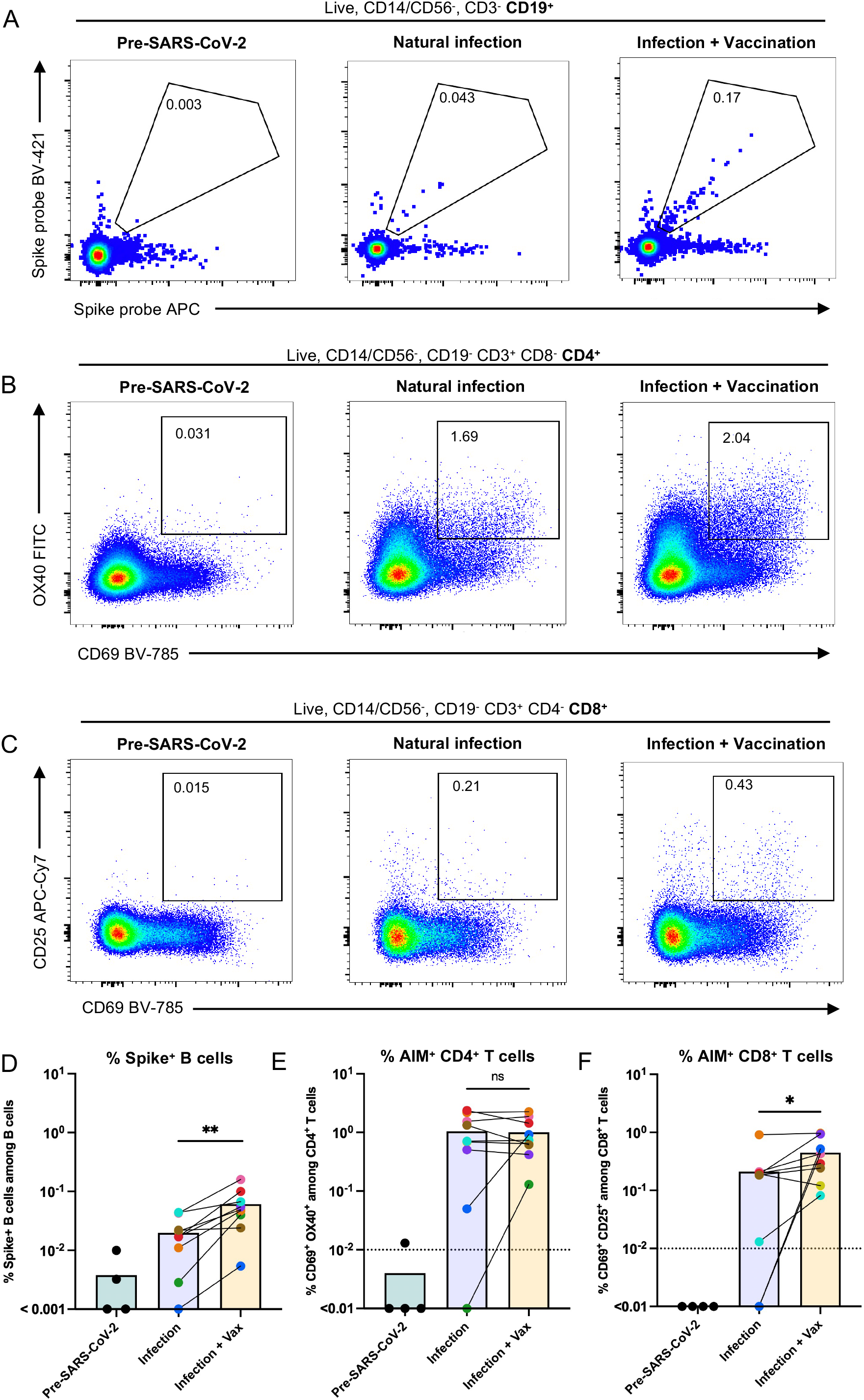
Spike-specific lymphocyte responses following natural infection and subsequent vaccination. A-C: Representative flow cytometry plots of SARS-CoV-2 spike protein-specific B cells (A), OX40/CD69^+^ CD4^+^ (B) and CD25/CD69^+^ CD8^+^ (C) T cells detected from PBMC using the BATTLE assay from a sample collected independently prior to the onset of the pandemic, and following natural infection and subsequent vaccination in the same individual (left to right, respectively). Pre-gating is shown above each panel. The frequency of cells within the double-positive gate among B cells, CD4+ T cells, and CD8+ T cells is shown inside the gates, respectively. D: The frequency among B cells of cells positive for both probes are shown, with each dot or dot pair representing an individual. Paired post-infection and post-infection plus vaccination samples from the same individual are connected by a solid line. Paired groups were analyzed by Wilcoxon matched-pairs signed rank test (n = 9, exact two-tailed p = 0.0039). E: The frequency among CD4^+^ T cells of cells positive for activation-induced markers OX40 and CD69 are shown, following subtraction of DMSO control values. Each dot or dot pair represents an individual. Paired post-infection and post-infection plus vaccination samples from the same individual are connected by a solid line. Paired groups were analyzed by Wilcoxon matched-pairs signed rank test (n = 9, exact two-tailed p > 0.9999). F: The frequency among CD8^+^T cells of cells positive for activation-induced markers CD25 and CD69 are shown, following subtraction of DMSO control values. Each dot or dot pair represents an individual. Paired post-infection and post-infection plus vaccination samples from the same individual are connected by a solid line. Paired groups were analyzed by Wilcoxon matched-pairs signed rank test (n = 9, exact two-tailed p = 0.0234).

Taken together, our data are in alignment with other studies in which an enhancement of spike-reactive adaptive B cell responses was observed in individuals who received SARS-CoV-2 mRNA vaccination following natural infection (Wang et al., 2021).

### SARS-CoV-2 mRNA vaccination differentially impacts the spike-specific peripheral CD4^+^ and CD8^+^ T cell compartments following natural infection

We next sought to determine whether spike-reactive CD4^+^ and CD8^+^ T cells within the same PBMC vial were similarly enhanced by mRNA vaccination after natural SARS-CoV-2 infection. We quantified the expression of activation-induced markers on these T cell subsets and found that, unlike for B cells targeting spike protein, the frequency of CD4^+^ spike peptide-activated T cells was not significantly enhanced following mRNA vaccination compared to infection alone (**Figure 3b, e**). Although significant changes were not observed for the full cohort, the frequency and number of AIM^+^ CD4^+^ T cells was increased following vaccination for some individuals, particularly for those with a lower initial spike-specific adaptive response. The same trend was observed for subjects with low AIM^+^ CD8^+^ T cell frequencies following natural infection, but unlike the spike-specific CD4^+^ T cell compartment, CD8^+^ T cells reactive to spike were marginally enhanced in frequency following mRNA vaccination (**Figure 3c, f**). The same patterns were present when adjusting for the number of cells used as input for the assay, although for AIM^+^ CD8^+^ T cells differences between the groups no longer reached statistical significance (**Supplementary Figure 3b, c**). As observed for spike-specific B cells, few AIM^+^ T cells were present in PBMC samples collected prior to the onset of the SARS-CoV-2 pandemic, and unlike T cells from exposed subjects, those from pre-SARS-CoV-2 samples were largely unresponsive to spike peptide pool stimulation (**Supplementary Figure 3d, e**).

Correlative analysis of pooled data from both infection and infection plus vaccination samples revealed a positive correlation between the frequency of SARS-CoV-2 spike-specific B cells and serum IgG titers against the spike RBD from the original Wuhan strain, as well as the alpha and beta variants (**Supplementary Figure 3f**). The frequency of AIM^+^ CD8^+^ T cells also positively correlated with anti-RBD IgG, but this was only significant for the original SARS-CoV-2 strain. These results propose a coordinated response to SARS-CoV-2 between B cells and CD8^+^ T cells, potentially without sustained synchronization with the CD4^+^ T cell compartment. Taken together, these data suggest that mRNA vaccination of previously-infected individuals influences T cell subsets asymmetrically, and may allow those who mount a sub-optimal response to infection to achieve an immunological setpoint not reached via natural infection.

## Discussion

Here we demonstrate that B cells and CD4^+^ and CD8^+^ T cells specific for the same antigen can be simultaneously identified and quantified in a single PBMC sample. Furthermore, our results suggest that the ability for SARS-CoV-2 spike-specific B and T cells to be enhanced by mRNA vaccination following natural SARS-CoV-2 infection may not be equal. These findings also imply that the incorporation of additional antigens to SARS-CoV-2 vaccines may improve the balance of SARS-CoV-2-elicited memory. Other studies have demonstrated that the spike-specific B and T cell responses are highly heterogeneous within the human population (Dan et al., 2021; Rodda et al., 2020; Sette and Crotty, 2021; Weiskopf et al., 2020), findings which are echoed here. Our work suggests that despite this heterogeneity, trends in the patterns of antigen-specific B and T cell expansion and contraction can be detected even within small cohorts. Future work will examine whether the same level of heterogeneity exists for phenotypic and transcriptomic measures within the same populations.

Although our study was limited to the examination of PBMC, it is possible that these observations extend to other sources of lymphocytes, such as lymph node aspirates, bronchioalveolar lavage fluid, and bone marrow. Antigen-specific lymphocytes have been identified within these specimens in humans and non-human primates (Corbett et al., 2021; Kaneko et al., 2020; Turner et al., 2021), suggesting that BATTLE can be applied to maximize the yield of information gathered from these valuable samples.

Unlike other studies, we did not detect substantial frequencies or numbers of SARS-CoV-2 spike-binding T cells in non-exposed individuals (Grifoni et al., 2020; Mateus et al., 2020). We attribute the disparity between our observations and others’ potentially to demographic variability between our sampling region and populations and the those sampled in other studies. It is also possible that the sensitivity of our assay was lower, although we consider this unlikely to be the case, as the quantities of antigen-specific T cells we observed post-infection and post-vaccination were well-aligned with those in the literature (Dan et al., 2021; Rydyznski Moderbacher et al., 2020).

The observation of increased quantities of spike-specific B cells following mRNA vaccination after natural infection supports the idea that mRNA vaccination is beneficial even after natural adaptive immunity has developed. Additionally, our results suggest that spike-reactive B cells may possess a capacity for vaccine-elicited expansion beyond that of spike-reactive CD4^+^ T cells. The mechanism by which this process may occur is still an active area of investigation, but an attractive hypothesis is that B and T cell populations each achieve an independent immunological setpoint following resolution of acute infection or vaccine-induced activation. As the epitopes recognized by B cells, CD4^+^ T cells, and CD8^+^ T cells, the timing and settings in which these events occur, and the environmental signals these population receive can differ considerably, it is tempting to speculate that uncoordinated dynamic expansion and contraction of B and T cells can lead to the generation of immunological memory in a compartmentalized manner. This interpretation would be supported with recent work describing coordinated and timely B and T cell responses that lead to the generation of superior adaptive immunity, and importantly that the lack thereof can have pathological consequences (Lucas et al., 2021; Painter et al., 2021; Zohar et al., 2020). BATTLE is an ideal approach that can be harnessed to explore these concepts.

Our study did not exclude the possibility that co-culture during T cell stimulation and/or BCR baiting has an impact on each cell population at the transcriptional level. Indeed, it will be important to examine the transcriptional signatures of T and B cells isolated via BATTLE compared to traditional assays to explore this possibility. If the co-stimulatory conditions do indeed cause activation or exhaustion of B or T cells, we envision the adaptation of this assay to study the outcome of any resulting cognate recognition-driven processes.

Given the paucity of antigen-specific memory lymphocytes following infection and vaccination, the findings here have the potential to increase the breadth of diagnostic and experimental protocols which rely on interrogation of these populations. Thus, our studies have potential implications beyond the study of SARS-CoV-2 and mRNA vaccination. We plan to adapt BATTLE to examine adaptive responses to additional antigens in the future.

## Acknowledgements

We gratefully acknowledge excellent technical assistance provided by the Upstate Medical University Flow Cytometry Core. We thank Lisa Phelps for her technical support. We thank Drs. J. Wilmore, H. Friberg, J. Currier, and G. Gromowski, and A. Wegman for their helpful comments.

The following reagents were obtained through BEI Resources, NIAID, NIH: Spike Glycoprotein (Stabilized) from SARS-Related Coronavirus 2, Wuhan-Hu-1 with C-Terminal Histidine and Avi Tags, Recombinant from HEK293F Cells, NR-53524, Peptide Array, SARS-Related Coronavirus 2 Spike (S) Glycoprotein, NR-52402, Spike Glycoprotein Receptor Binding Domain (RBD) from SARS-Related Coronavirus 2, United Kingdom Variant with C-Terminal Histidine Tag, Recombinant from HEK293 Cells, NR-54004, and Spike Glycoprotein Receptor Binding Domain (RBD) from SARS Related Coronavirus 2, Beta Variant with C-Terminal Histidine Tag, Recombinant from HEK293 Cells, NR-55278.

## Contributions

K.L.N. conceptualized and executed experiments, analyzed the data, constructed figures and wrote and edited the manuscript. M.W. aided in execution and analysis of serum ELISAs. S.J.T. and T. P.E. contributed to project administration and funding acquisition. A.T.W. conceptualized experiments, and aided in the analysis of data, project administration, funding acquisition, and the writing and review of the manuscript. All authors participated in final review of the manuscript.

## Data Availability

Flow cytometry data has been submitted under ID FR-FCM-Z4PB and annotated at http://flowrepository.org. All other data is available upon request.

---

**Supplementary Figure 1: BATTLE assay gating strategy.**

The flow cytometry gating strategy used to identify SARS-CoV-2 spike protein-specific B and T cells is shown using a representative sample.

**Supplementary Figure 2: SARS-CoV-2 variant spike RBD serology.**

A-D: Paired sera following natural infection and after subsequent vaccination from the same individuals were assessed for IgM and IgG binding to SARS-CoV-2 alpha (A, B) or beta (C, D) variant spike RBD proteins. Each dot color represents an individual and pairs are connected by a solid line. Differences between infection and infection + vaccination groups were analyzed by Wilcoxon matched-pairs signed rank test (n = 9, exact two-tailed p = 0.6250, 0.0195, 0.3125, 0.0469 (A-D)).

**Supplementary Figure 3: AIM^+^ T cell analysis and correlative analysis.**

A: The number of B cells positive for both probes per 1 x 10^6^ cells plated for stimulation is shown, with each dot or dot pair representing an individual. Paired post-infection and post-infection plus vaccination samples from the same individual are connected by a solid line. Paired groups were analyzed by Wilcoxon matched-pairs signed rank test (n = 9, exact two-tailed p = 0.0039).

B: The number of CD4^+^ T cells positive for activation-induced markers OX40 and CD69 per 1 x 10^6^ cells plated for stimulation is shown, following subtraction of DMSO control values. Each dot or dot pair represents an individual. Paired post-infection and post-infection plus vaccination samples from the same individual are connected by a solid line. Paired groups were analyzed by Wilcoxon matched-pairs signed rank test (n = 9, exact two-tailed p = 0.5703).

C: The number of CD8^+^ T cells of cells positive for activation-induced markers CD25 and CD69 are shown, following subtraction of DMSO control values. Each dot or dot pair represents an individual. Paired post-infection and post-infection plus vaccination samples from the same individual are connected by a solid line. Paired groups were analyzed by Wilcoxon matched-pairs signed rank test (n = 9, exact two-tailed p = 0.2500).

D. The frequency of AIM^+^ CD4^+^ T cells among total CD4^+^ T cells is shown with each pair of dots representing an individual. Paired DMSO control and peptide pool-stimulated samples from the same individual are connected by a solid line (n = 4 pre-SARS-CoV-2, 20 post-SARS-CoV-2). Subjects exposed to SARS-CoV-2 mRNA vaccination only (turquoise), natural infection only (purple), and infection plus vaccination (orange) are shown.

E. The frequency of AIM^+^ CD8^+^ T cells among total CD8^+^ T cells is shown with each pair of dots representing an individual. Paired DMSO control and peptide pool-stimulated samples from the same individual are connected by a solid line (n = 4 pre-SARS-CoV-2, 20 post-SARS-CoV-2). Subjects exposed to SARS-CoV-2 mRNA vaccination only (turquoise), natural infection only (purple), and infection plus vaccination (orange) are shown.

F. Spearman’s rho rank correlation coefficients are graphed by circle color according to the scale on the right of the panel. The size of the circle indicates increasing statistical significance. Correlation matrix was generated from data as presented in the previous figures representing the combined SARS-CoV-2 infection and vaccination-elicited immune response.

Supplementary Table 1: Fluorescent antibodies used in BATTLE assay.

