## Supplementary data for "Simultaneous analysis of antigen-specific B and T cells after SARS-CoV-2 infection and vaccination"

**Supplementary Fig. 1 Gating strategy**

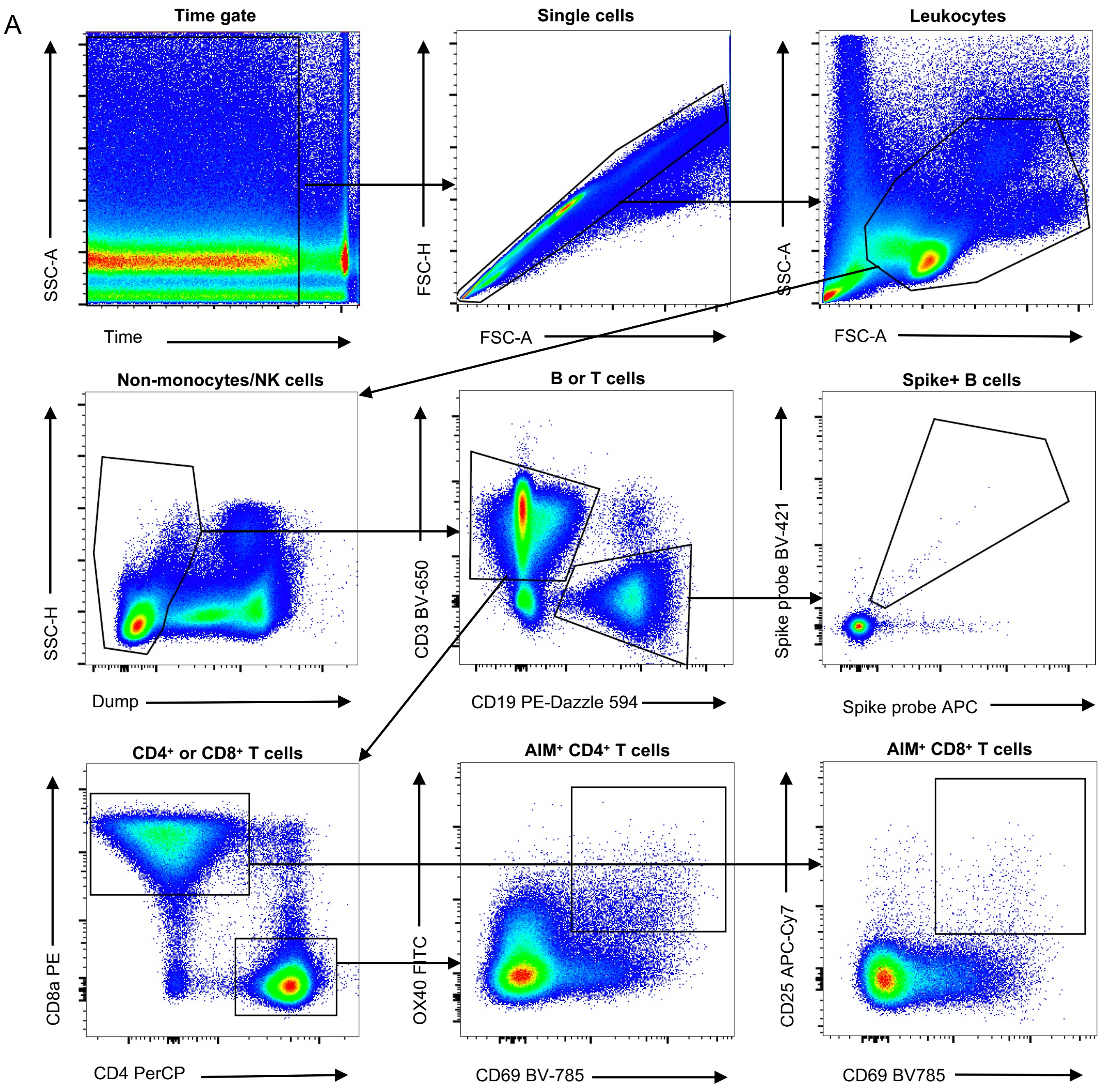

**Supplementary Fig. 2: SARS-CoV-2 variant spike RBD serology**

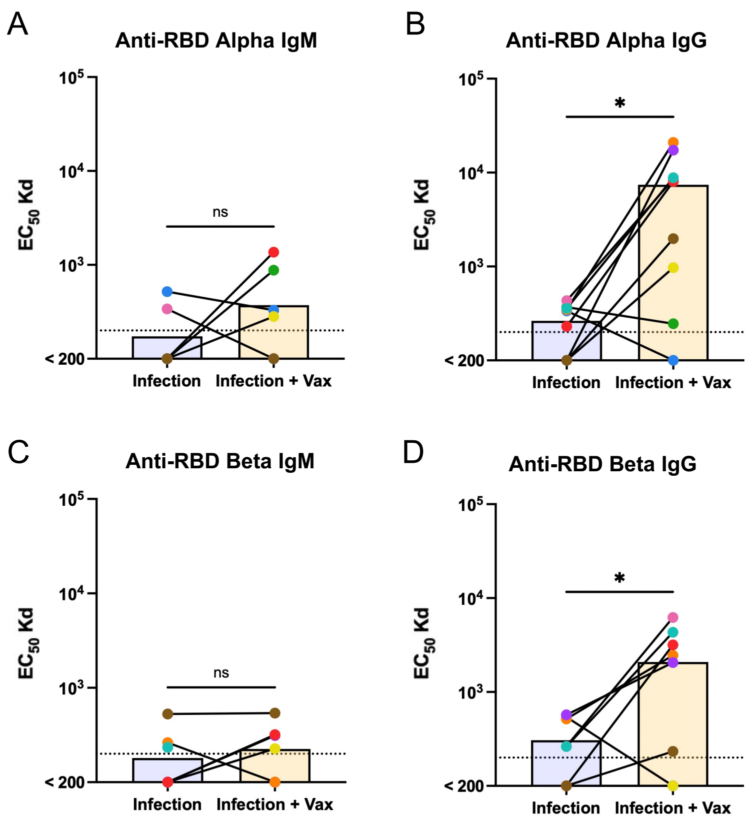

**Supplementary Fig. 3: AIM$^+$ T cell analysis and correlations**

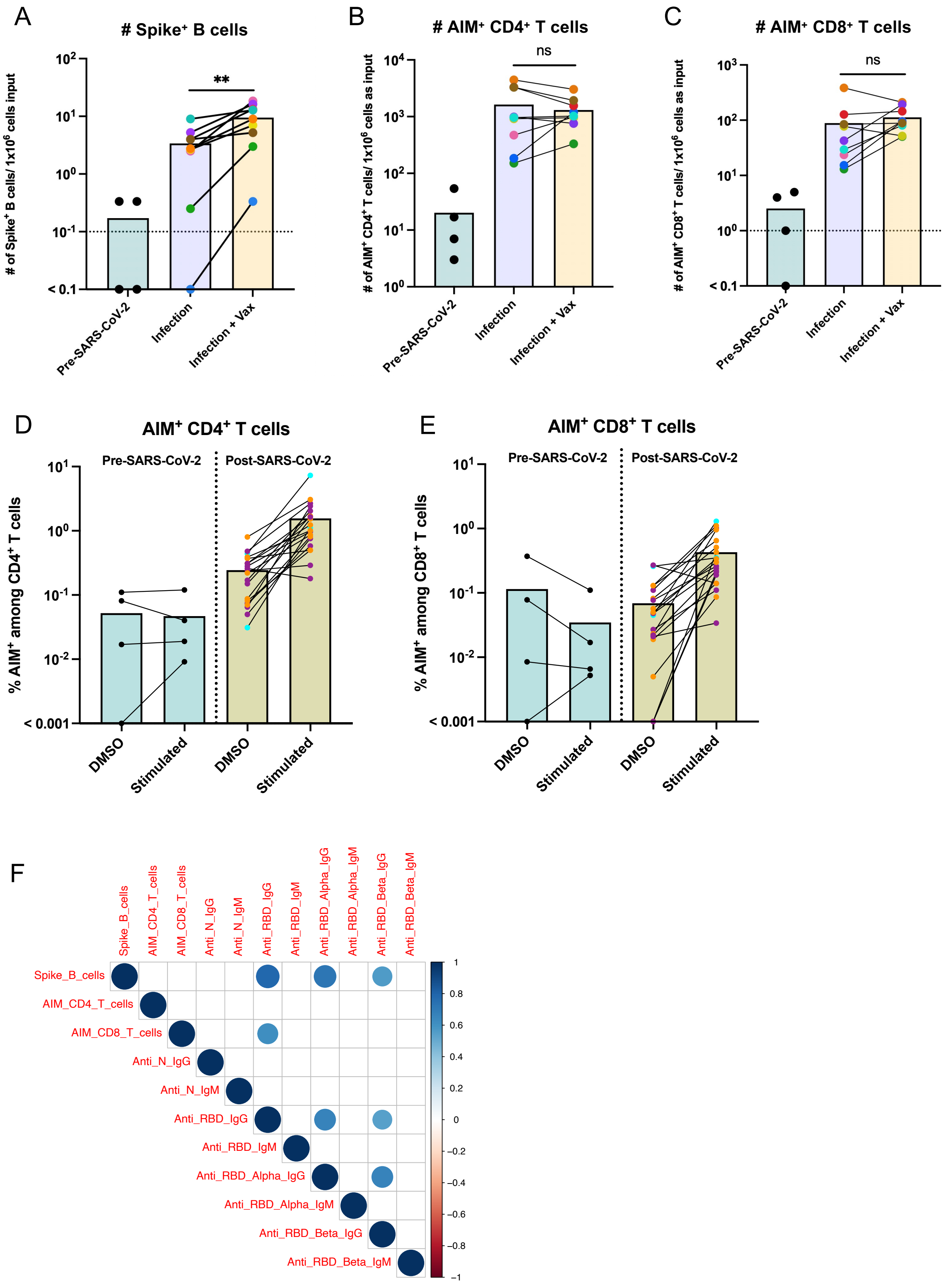

**Supplementary Table 1**

| ANTIBODY | SOURCE | IDENTIFIER |
|---|---|---|
| FITC Mouse anti-human CD134 (OX40) | BioLegend | Cat. 350006 |
| PE-Dazzle 594 Mouse anti-human CD19 | BioLegend | Cat. 302251 |
| PE Mouse anti-human CD8a | BioLegend | Cat. 301007 |
| PerCP Mouse anti-human CD4 | BioLegend | Cat. 317431 |
| APC anti-human streptavidin | BD Biosciences | Cat. 554067 |
| APC-Cy7 Mouse anti-human CD25 | BioLegend | Cat. 302613 |
| BV421 anti-human streptavidin | BD Biosciences | Cat. 563259 |
| BV510 Mouse anti-human CD14 | BioLegend | Cat. 301841 |
| BV510 Mouse anti-human CD56 | BioLegend | Cat. 318339 |
| BV650 Mouse anti-human CD3 | BioLegend | Cat. 317323 |
| BV785 Mouse anti-human CD69 | BioLegend | Cat. 310931 |
| Biotin anti-human CD19 | BioLegend | Cat. 302203 |
